# Artificial mosaic brain evolution of relative telencephalon size improves cognitive performance in the guppy (*Poecilia reticulata*)

**DOI:** 10.1101/2021.04.03.438307

**Authors:** Zegni Triki, Stephanie Fong, Mirjam Amcoff, Niclas Kolm

**Author notes:** **Corresponding author:** Zegni Triki.

## Abstract

The telencephalon is a brain region believed to have played an essential role during cognitive evolution in vertebrates. However, till now, all the evidence on the evolutionary association between telencephalon size and cognition stem from comparative studies. To experimentally investigate the potential evolutionary association between cognitive abilities and telencephalon size, we used male guppies artificially selected for large and small telencephalon relative to the rest of the brain. In a detour task, we tested a functionally important aspect of executive cognitive ability; inhibitory control abilities. We found that males with larger telencephalon outperformed males with smaller telencephalon. They showed faster improvement in performance during detour training and were more successful in reaching the food reward without touching the transparent barrier. Together, our findings provide the first experimental evidence showing that evolutionary enlargements of relative telencephalon size confer cognitive benefits, supporting an important role for mosaic brain evolution during cognitive evolution.

## Introduction

Cognition is the ability to acquire, process, retain information and act on it (Shettleworth, 2010). Variation in cognition is extensive at all taxonomic levels (Shettleworth, 2010; MacLean et al., 2014; Benson-Amram et al., 2016), and often surprisingly large even across individuals in the same population (Thornton & Lukas, 2012; Ashton et al., 2018; Boogert et al., 2018; Triki et al., 2019b). But an important question remains largely unanswered (e.g. Thornton & Lukas, 2012): how does cognition evolve?

Till now, virtually all existing empirical work on cognitive evolution is based on comparative studies of the link between various aspects of brain morphology and cognitive abilities ((Jerison, 1973; Timmermans et al., 2000; Deaner et al., 2007; Reader, Hager & Laland, 2011; MacLean et al., 2014), see also (Kotrschal et al., 2013; Benson-Amram et al., 2016; Buechel et al., 2018)). One of the most important hypotheses for cognitive evolution in this context is the mosaic brain evolution hypothesis (Barton & Harvey, 2000). This hypothesis suggests that direct selection on individual cognitive abilities leads to independent changes in the size of key brain regions with distinct functional properties (Barton & Harvey, 2000; Striedter, 2005). Ecological demands from the environment can thus generate selection on specific parts of the brain to achieve higher cognitive capacities to cope with such challenges (Kolm et al., 2009; Reader, Hager & Laland, 2011; Ebbesson & Braithwaite, 2012; van Schaik, Isler & Burkart, 2012; Triki et al., 2019a). The telencephalon is a brain region of particular interest in studies of cognitive evolution and mosaic brain evolution. The reason for this is because it controls key cognitive functions such as memory, spatial memory, fear conditioning, and decision making (Salas et al., 1996; López et al., 2000; Portavella et al., 2002; O’Connell & Hofmann, 2011). Phylogenetic comparative analyses on vertebrate telencephalon evolution are not uncommon. For instance, the neocortex makes up the majority of the telencephalon in primates (Briscoe & Ragsdale, 2019). Group social complexity, diet and cognitive performance are positively associated with neocortex size in this taxon (Shultz & Dunbar, 2010; Reader, Hager & Laland, 2011; DeCasien & Higham, 2019). In birds, larger telencephalon size is associated with cognitive capacity in the form of feeding innovation (Timmermans et al., 2000). In fish, relative telencephalon size is associated with habitat complexity (Gonzalez-Voyer & Kolm, 2010; White & Brown, 2015), and with sex-specific differences in behaviour and home range size (Kolm et al., 2009; Costa et al., 2011). Fish forebrain (i.e., telencephalon and diencephalon) size is also associated with the capacity to optimise decision-rules as a function of the social environment (Triki et al., 2020), known as social competence (Taborsky & Oliveira, 2012). These reports of increases in the telencephalon’s proportional size in relation to cognitively demanding behaviour support general implications from the mosaic brain evolution hypothesis. But experimental data is needed to establish whether mosaic brain evolution can be an important engine of cognitive evolution.

A recent artificial selection experiment in the guppy (*Poecilia reticulata*), changing the size of the telencephalon relative to the rest of the brain (Fong et al., 2021, preprint) provides experimental support for that separate brain regions can evolve independently (Barton & Harvey, 2000). These selection lines now offer a rare opportunity to experimentally test whether evolutionary telencephalon enlargement is accompanied by enhanced cognitive abilities, consistent with mosaic brain evolution driving cognitive evolution. After three generations of selection, divergence in telencephalon size ratio has reached 5 % in males and 7 % in females without apparent changes in the other brain regions between the two selection lines (Fong et al., 2021, preprint). Importantly, the fish telencephalon has been suggested to share at least some of the functions of the mammalian neocortex and the bird telencephalon. Invasive methods have shown that individuals with lesioned or ablated telencephalon had impaired spatial memory (Salas et al., 1996; Portavella et al., 2002), and deficiencies in learning abilities (Portavella et al., 2002). Moreover, the telencephalon in fish also shares conserved gene expression networks with the mammalian neocortex (O’Connell & Hofmann 2011, 2012). Hence, targeting this region for artificial selection should be suitable for the study of cognitive evolution also from a general perspective.

In this study, we tested male guppies from the telencephalon selection lines for their cognitive abilities in a detour task. This task has been performed on several species across taxa (fish, birds, and mammals), and it tests for the executive function “self-control” (Boogert et al., 2011; MacLean et al., 2014; Fagnani et al., 2016; Lucon-Xiccato, Gatto & Bisazza, 2017; Brandão, Fernandes & Gonçalves-de-Freitas, 2019). Executive functions (i.e., self-control, working memory, and cognitive flexibility) are often referred to as control mechanisms of general-purpose that modulate different cognitive subprocesses (Miyake et al., 2000). The self-control executive function requires inhibitory control abilities to override motor impulses, resulting in adaptive goal-oriented behaviours when correctly performed (Kabadayi, Bobrowicz & Osvath, 2018). Following the logic of more neural tissue conferring “better” cognitive abilities, we expected that males from the up-selected lines, with larger telencephalon, should show higher inhibitory control abilities than males from the down-selected lines with small telencephalon. We also measured the relative telencephalon size in the tested fish to verify that the expected divergence in telencephalon size between the selection lines was evident in the sampled fish.

## Results

Performance, both during the pre-training and test phases of the detour task, improved significantly with additional trials (*p* < 0.001, Table1, Fig. 1 & Fig. 2). In the pre-training phase, we found a significant difference between up’ and down-selected telencephalon lines in the speed of learning the task (i.e., number of time-bins to reach the food reward; See Methods) over trials (*p* = 0.016, Fig. 1). Up-selected lines had a significantly steeper performance-trial slope (*post hoc* test: emtrend = −0.349, *p* < 0.001) than the down-selected lines (emtrend = −0.138, *p* = 0.019). In the test phase, latency to reach the food reward was not significantly affected by the telencephalon size (Fig. 2a, Table 1). Reaching the food reward without touching the barrier is a crucial measurement of inhibitory control ability (see review by Kabadayi, Bobrowicz & Osvath, 2018). Our results showed that up-selected lines outperformed the down-selected lines in detouring without touching the barrier (*p* = 0.013, Fig. 2b).

**Figure 1.**
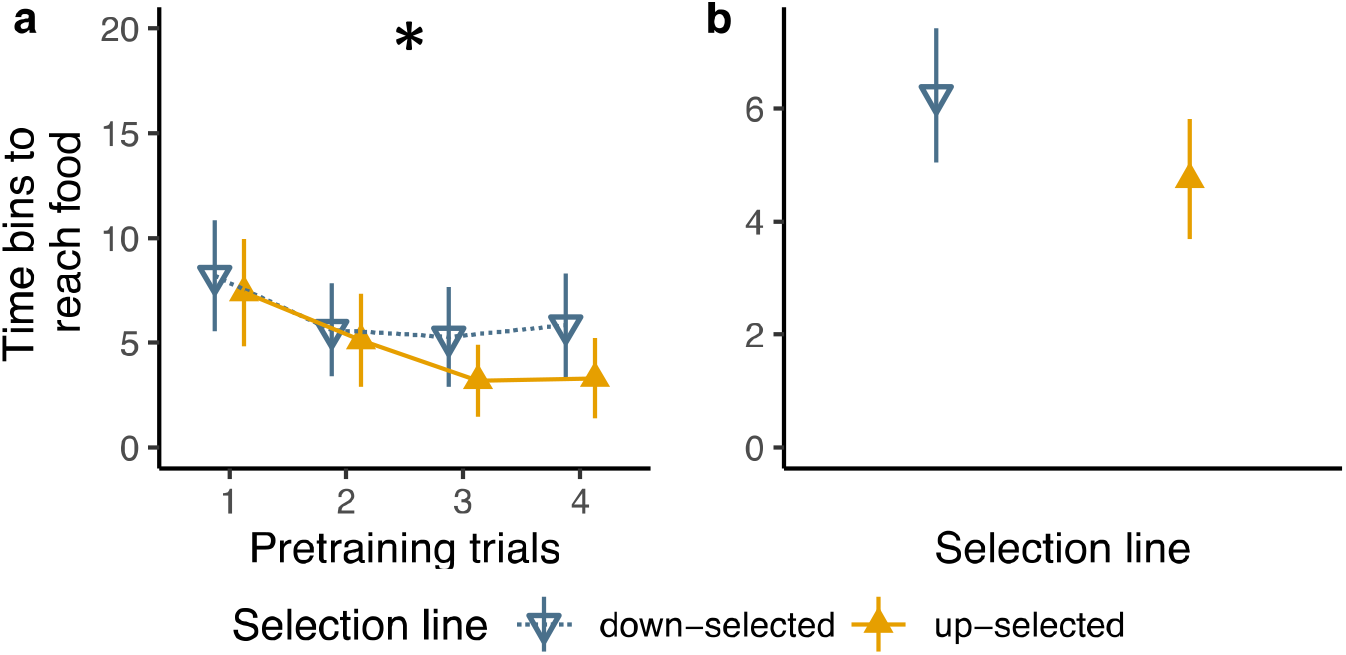
Performance in the pre-training trials. Mean and 95 % confidence levels of the number of time-bins elapsed until fish reached the food reward: (**a**) per trial and selection line, and (**b**) for all trials per selection line. Values are estimated from 36 males from the up selected lines and 36 males from the down-selected lines in the pre-training phase. GLMM: * significant interaction between selection treatment and trials, *p <*0.05.

**Figure 2.**
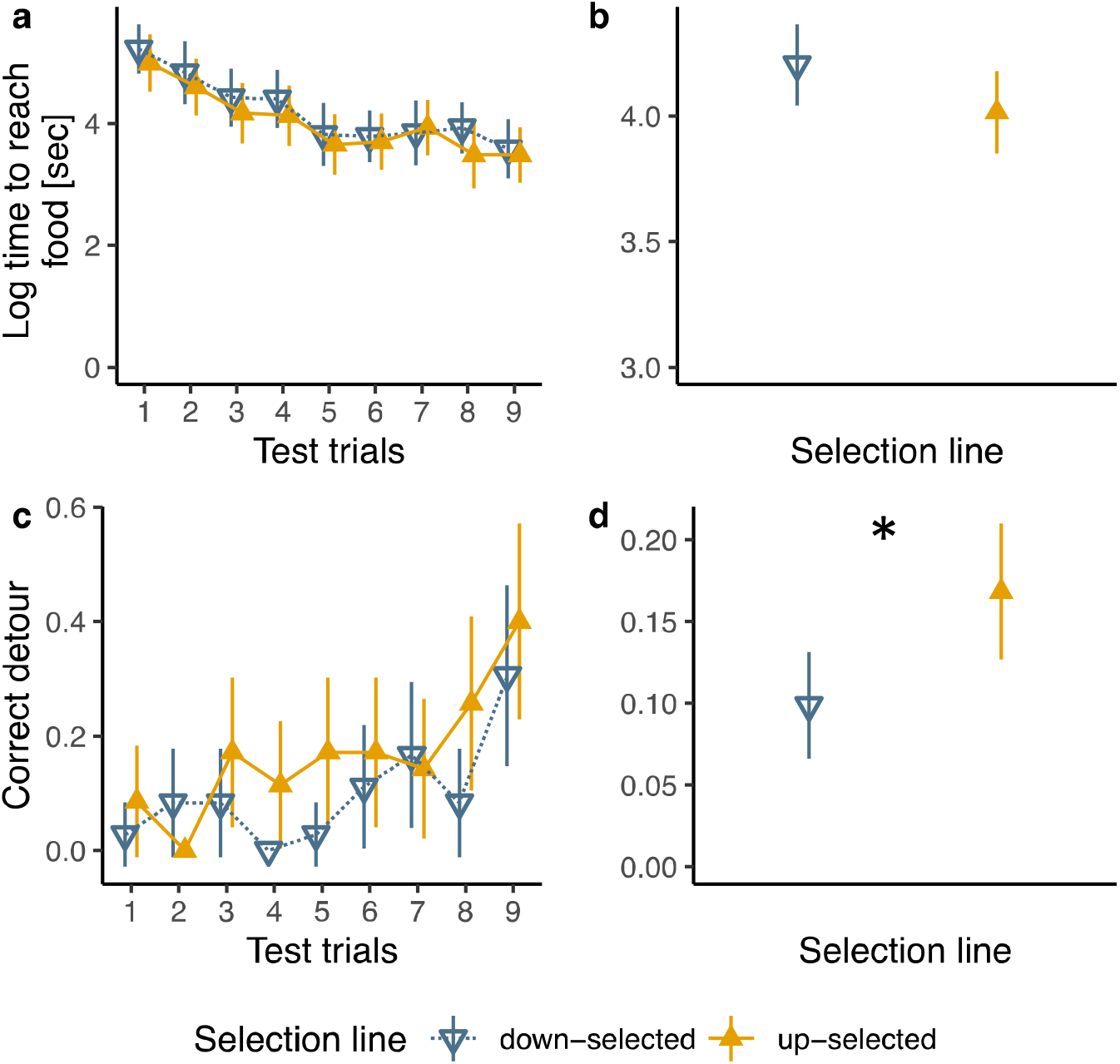
Performance in the test trials. Mean and 95 % confidence level of time to reach a food reward: (**a**) per trial and selection line, and (**b**) for all trials per selection line. Mean and 95 % confidence level of correct detours (i.e., detouring the transparent cup without touching it): (**c**) per trial and selection line, and (**d**) for all trials per selection line. Values are estimated from 35 males from the up-selected lines and 36 males from the down-selected lines in the test phase. GLMM: * significant effect of selection treatment, *p <*0.05.

**Figure 3.**
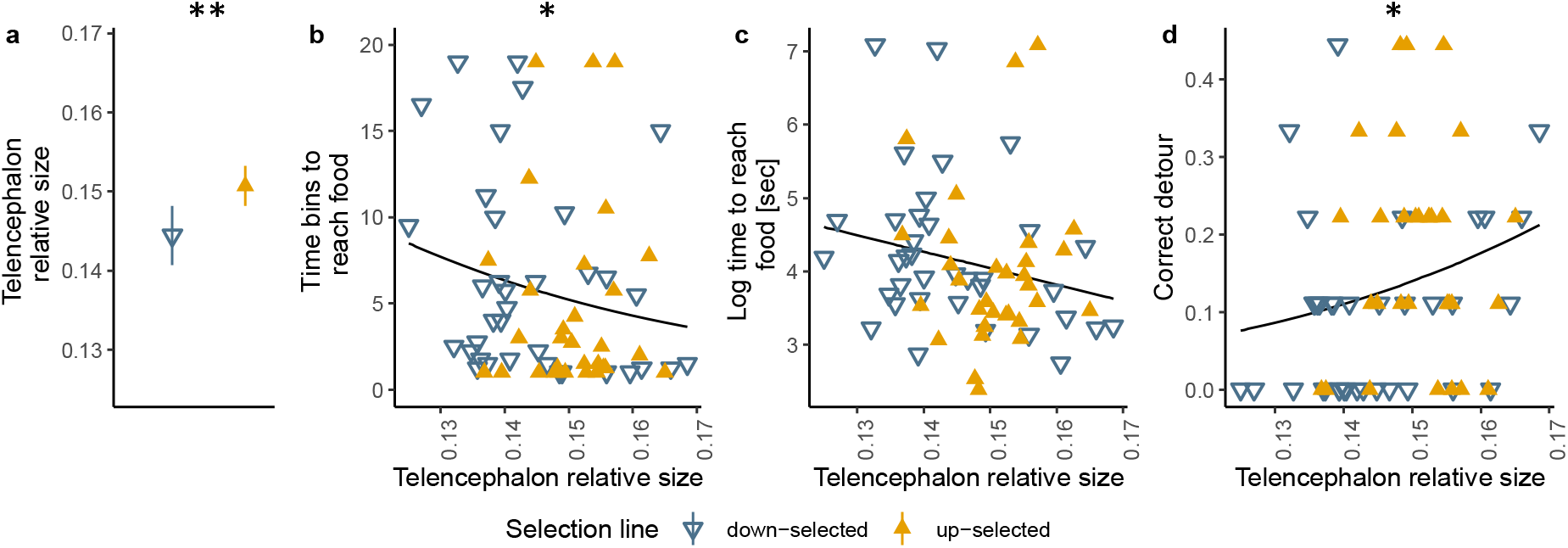
Individual telencephalon relative size, selection lines and performance. (**a**) Mean and 95 % confidence level of telencephalon size proportion from 31 up’ and 36 down selected male guppies. Regression relationship of telencephalon size proportion and (**b**) time bins to reach food reward in the pre-training trials, (**c**) time elapsed to reach the food reward, (**d**) correct detours. The filled (31 up-selected) and open triangles (36 down-selected) are the average performance of every individual over trials. GLMM: \**p <*0.05; ** *p < 0.01.*

**Table 1.** Summary table of the statistical outcomes from models testing how far cognitive performance in the detour task is predicted by either selection line or individual telencephalon size proportion. Significant results with an *alpha* set at ≤ 0.05 are indicated in bold font.

| Performance and selection line | Pre-training phase. Number of bins to reach food reward (Negative binomial distribution) |  |  | Test phase. Time to detour (Gaussian distribution) |  |  | Test phase. Correct detour (Binomial distribution) |  |  |
| --- | --- | --- | --- | --- | --- | --- | --- | --- | --- |
| Predicting variables | $Chi^2$ | $df$ | $p$-value | $Chi^2$ | $df$ | $p$-value | $Chi^2$ | $df$ | $p$-value |
| Selection line | 2.559 | 1 | 0.109 | 0.662 | 1 | 0.416 | **6.120** | **1** | **0.013** |
| Trial | **28.863** | **1** | **< 0.001** | **124.539** | **1** | **< 0.001** | **28.357** | **1** | **< 0.001** |
| Selection treatment x Trial | **5.841** | **1** | **0.016** | 0.155 | 1 | 0.694 | 0.050 | 1 | 0.822 |
| Replicate | 0.974 | 2 | 0.614 | **6.722** | **2** | **0.035** | **8.105** | **2** | **0.017** |
| Marginal $R^2$ | **0.10** | | | **0.14** | | | **0.19** | | |
| Conditional $R^2$ | **0.68** | | | **0.51** | | | **0.23** | | |
| Sample size | N = 36 up-selected<br>N = 36 down-selected |  |  | N = 35 up-selected<br>N = 36 down-selected |  |  | N = 35 up-selected<br>N = 36 down-selected |  |  |
| Performance and telencephalon size proportion | Pre-training phase. Number of bins to reach food reward (Negative binomial distribution) |  |  | Test phase. Time to detour (Gaussian distribution) |  |  | Test phase. Correct detour (Binomial distribution) |  |  |
| Predicting variables | $Chi^2$ | $df$ | $p$-value | $Chi^2$ | $df$ | $p$-value | $Chi^2$ | $df$ | $p$-value |
| Telencephalon size proportion | **3.958** | **1** | **0.046** | 3.133 | 1 | 0.076 | **4.139** | **1** | **0.042** |
| Trial | **23.542** | **1** | **< 0.001** | **106.698** | **1** | **< 0.001** | **24.350** | **1** | **< 0.001** |
| Telencephalon size proportion x Trial | 0.769 | 1 | 0.380 | 0.001 | 1 | 0.973 | 0.048 | 1 | 0.826 |
| Marginal $R^2$ | **0.09** | | | **0.11** | | | **0.13** | | |
| Conditional $R^2$ | **0.62** | | | **0.51** | | | **0.21** | | |
| Sample size | N = 31 up-selected<br>N = 36 down-selected |  |  | N = 31 up-selected<br>N = 36 down-selected |  |  | N = 31 up-selected<br>N = 36 down-selected |  |  |
| Marginal $R^2$ refers to explained variance by fixed factors, while conditional $R^2$ refers to explained variance by both fixed and random factors. | | | | | | | | | |

From measurements of relative telencephalon size, as the size proportion of total brain size, we found that the tested fish differed significantly in telencephalon size between the up’ and down-selected lines, a difference of 4.9 % (LM: N = 31 up-selected vs N = 36 down-selected, F-value = 8.38, *p* = 0.005, adjusted-R^2^ = 0.12, Fig. 4a). In support of the tests employing selection line as a predictor, correlation analyses between telencephalon size proportion and performance in the detour task showed similar outcomes (Fig. 4b-d). Telencephalon size co-varied negatively with number of time-bins required to reach the food reward during the pre-training phase, whereas it co-varied positively with proportion of correct detours in the test phase. In sum, as the telencephalon proportion gets larger, fish detour performance improved both during the pre-training and during the test phase (Table 1).

## Discussion

The use of artificial selection lines with differences in relative telencephalon size provides the first experimental evidence that evolutionary changes in the size of this region affect cognitive abilities in an important executive function.

The most relevant finding linked to our prediction - that a larger telencephalon would confer cognitive benefits - is that animals from the up-selected lines improved faster during the pre-training phase, and outperformed the down-selected lines in the test phase. This finding was mirrored by a positive relationship between the individual relative telencephalon size of every tested fish and performance in the detour task. Overall, we found significant improvement over trials in both the pre-training and test phases, suggesting a learning process in the test paradigm through repeated testing (Lucon-Xiccato, Gatto & Bisazza, 2017; Kabadayi, Bobrowicz & Osvath, 2018). Several cognitive processes might be responsible for this difference in improvement, such as differences in either spatial navigation, learning, working memory, or route planning (reviewed in Kabadayi, Bobrowicz & Osvath, 2018). Interestingly, in primates, the proximate neurological underpinnings of inhibitory control abilities are often attributed to the prefrontal cortex and its analogous structures in the avian telencephalon (reviewed in Kabadayi, Bobrowicz & Osvath, 2018). Our results suggest a similar role for telencephalon in the guppy telencephalon size selection lines, again supporting a highly general functional role of the telencephalon across distant taxonomic groups (Striedter, 2005; Striedter & Northcutt, 2019). Regardless of the exact mechanism behind differences in performance, our results suggest that strong directional selection on relative telencephalon size can lead to improved cognitive abilities. This is an important finding because it indicates that cognition can evolve rapidly at the within-species level via targeted changes in specific brain regions. This, in turn, provides direct support for the mosaic brain evolution hypothesis, the idea that relatively independent evolutionary changes in brain region sizes and functions are important for cognitive evolution (Barton & Harvey, 2000). The difference in relative telencephalon size between third generation up’ and down-selected telencephalon lines used here was approximately 5 % (Fong et al., 2021, preprint), which was confirmed in the subsample of fish tested in our study (4.9 % difference). That such a small difference in this region yielded significant differences in cognitive performance suggests a disproportionate relationship between key brain region size and function (Finlay, Darlington & Nicastro, 2001). This highlights the need to consider also small differences in relative brain region size when investigating behavioural differences. Additional cognitive tasks tackling different cognitive processes, as well as cognitive tests in females, are needed to increase our understanding of the cognitive function of a larger telencephalon in these selection lines. Although it was already established that there is no difference in reproductive output between the up’ and down-selected telencephalon lines (Fong et al., 2021, preprint), additional work is also needed to quantify the potential energetic costs associated with evolutionary changes in brain morphology and cognitive ability.

Self-inhibition is a general aspect of guppy cognition and thus highly ecologically relevant. For instance, another recent study found that female guppies can show inhibitory control abilities (Lucon-Xiccato, Gatto & Bisazza, 2017). Both this study and another taxonomically broader study on primates and birds (MacLean et al., 2014) used an opaque barrier to pre-train individuals on detouring. Animals in these studies scored between 26 % and 100 % correct detours, with guppies having a score of 58 % (MacLean et al., 2014; Lucon-Xiccato, Gatto & Bisazza, 2017). In the present study, we observed a relatively low detour performance with the down’ and up-selected telencephalon scoring on average 10 % and 17% correct detours, respectively. It is possible that first training individuals with an opaque barrier improves performance (Santos, Ericson & Hauser, 1999; Juszczak & Miller, 2016). We opted for a more challenging detour test paradigm, where we used a transparent barrier throughout the entire experiment. The visibility of the target without pre-training with an opaque barrier may make the task more challenging, for instance, because of increased conflict between the visual stimulus and tactical perception (see Juszczak & Miller, 2016). Indeed, familiarisation with the test increased throughout the experiment, with performance improving to a score of 30 % and 40 % correct detours in the last trial for down’ and up-selected telencephalon lines respectively. Hence, despite improvement during the trials, the average gap in performance between the down’ and up-selected telencephalon lines remained constant.

## Conclusions

We show that artificial selection mimicking evolutionary increases and decreases in telencephalon size leads to improved inhibitory control abilities. The demonstrated link between rapid evolutionary changes in brain morphology and cognitive abilities provides new insights into how cognitive abilities can evolve, and provides support for that mosaic brain evolution can be an important mechanism behind individual variation in brain morphology and cognition.

## Methods

### Study animals

We used three replicated laboratory lines of Trinidadian guppies (*Poecilia reticulata*) artificially selected for having relatively large or small telencephalon (Fong et al., 2021, preprint). Briefly, the artificial selection was made such that 225 breeding pairs (F_0_) were set up and allowed to produce at least two clutches. Then the brains of the parents were dissected, and the volumes of five major brain parts (telencephalon, optic tectum, hypothalamus, cerebellum and dorsal medulla) and the olfactory bulb were measured using the methods by Pollen et al. (2007), and Gonzalez-Voyer et al. (2009). Based on these volume measurements, new breeding pairs were ranked using the sum of residuals for relative telencephalon size in relation to the rest of the brain. Afterwards, offspring from the higher-rank and lower-rank parent pairs were set up to produce the next generation (F_1_), which resulted in up’ and down-selected telencephalon lines. The process was repeated for successive generations. In the present study, we used male guppies from the up’ and down-selected populations from the third generation of selection (F_3_) with a 5 % average difference in telencephalon size (Fong et al., 2021, preprint). We used only males in the present study because we did not aim to test sex differences in cognitive performance.

Adult male guppies from each of the three replicates of up’ and down-selected telencephalon lines were kept in groups of 40 individuals in glass tanks (length × width × height; 60 × 22 × 27 cm) with aerated water. The room had an ambient temperature of ~ 26 °C with a light schedule of 12 hours light and 12 hours dark. We fed the guppies *ad libitum* with fish flakes and newly hatched brine shrimp six days per week.

### Detour task

Data were collected between January and February 2020 in the Zoology department aquarium facilities at Stockholm University. To test guppies from up’ and down-selected telencephalon lines in the detour task, we housed 72 male guppies across the three replicates (N = 12 fish per replicate and selection treatment) in individual experimental aquaria. Every experimental aquarium (40 × 15 × 15 cm) had 2 cm gravel, an artificial plant as enrichment, and continuously aerated water. Two sliding doors, one transparent and one opaque, divided each aquarium in two compartments: a housing and test compartment. The experimental aquaria were aligned next to each other and visibility was allowed between the housing compartments but not test compartments. This helped to reduce potential stress in the fish due to isolation, as well as biases in the performance (e.g. from social learning). Another member of the lab netted out the experimental fish and marked the experimental aquaria with running numbers, thus ensuring the experimenter (ZT) was blind to the telencephalon selection line treatment of the individual fish.

#### Acclimation

We first allowed fish to acclimate for ten days in the experimental aquaria before any trials. On the fifth day of acclimation, we started habituating fish twice a day to eat a defrosted adult brine shrimp off a green spot drawn on a white acrylic plate placed at the bottom of the test compartment.

#### Pre-training phase

On the 11th day, we started inserting a see-through plastic cup (5 cm length and 4 cm diameter), with the opening facing up (loosely adapted from Wallis et al., 2001). The cup was placed on top of the green feeding spot. Thus, we could place a food item (a defrosted adult brine shrimp) inside the cup but still on the top of the visible green feeding spot (see Supplementary Video S1). Once the cup with food was set, we pulled up the opaque sliding door followed by the see-through door, which gave the fish a few seconds to see and assess the set-up before having access to the test compartment. To reach the food item, fish had to detour the physical barrier (i.e., the plastic cup walls) and swim inside the cup aiming for the food placed on the green spot. We provided a maximum of 380 minutes per trial to reach the food item with one trial a day for four consecutive days. Here, the individual performance consisted of the number of “bins” until reaching the food; that is, the 380 minutes were divided into 19 time-bins of 20 minutes each per day.

#### Test phase

We provided a two-day break between the pre-training and test phase. Then, over five days of testing, we ran one test trial on the first day and two test trials per day for the following four days. Every trial consisted of a similar paradigm as in the pre-training phase. The only difference was that fish had a maximum of 20 minutes per trial to reach the food reward. Here, we assayed performance by recording the time in seconds, starting from when we pulled up the see-through door until the fish reached the food item. We also observed whether fish detoured correctly (MacLean et al., 2014; Lucon-Xiccato, Gatto & Bisazza, 2017; Kabadayi, Bobrowicz & Osvath, 2018), i.e., whether they detoured without touching the cup (for an example, see Supplementary Video S1). Five fish died during the experiment: one fish died after the pre-training phase, while the other four fish died after the experiment. The cause of death was that fish jumped out of the aquaria overnight. All five fish were from the up-selected lines.

### Volume measurements of major brain regions

After completing the detour task, we euthanised the fish with an overdose of benzocaine (0.4 g/1), and fixed their bodies with 4 % paraformaldehyde in phosphate-buffered saline (PBS) for five days. We then performed two washing steps in PBS for 10 min each before storage at 4 °C pending brain dissection. We measured fish standard length (SL) to the nearest 0.01 millimetre using digital callipers (mean ± SD: up-selected lines 17.27 ± 0.77, down-selected lines 17.69 ± 0.62 mm). Afterwards, we dissected fish brains out of the skull, and took pictures from the dorsal, right lateral, left lateral and ventral view of the brains under a stereo zoom microscope Leica MZFLIII^®^ with a digital camera Leica DFC 490. Importantly, we remained blind to the treatment of the fish during brain measurements. Using ImageJ open software, these images were used to estimate the length (*L*), width (*W*) and height (*H*) of the telencephalon, optic tectum, cerebellum, dorsal medulla, hypothalamus and olfactory bulb. By fitting *L*, *W*, and *H* in an ellipsoid function (1) (based on Pollen et al., 2007; and White & Brown, 2015), we calculated the volume (*V*) of every brain part (in mm^3^):

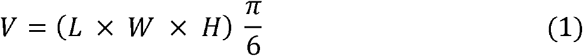

Finally, to estimate telencephalon relative size, we computed the size proportion of the telencephalon part from the total brain volume (i.e., the sum of all brain regions volume).

### Statistical analyses

For the statistical analyses and figures, we replaced missing values from individuals who failed to reach the food reward within the maximum allowed time of 19 bins and 20 minutes in the pre-training and test phase, respectively. To do so, we attributed a fixed value of 19 bins for data from the pre-training phase and a value of 20 minutes for the data from the test phase. Out of 288 data points from the pre-training phase, there were 31 failures from 16 down-selected fish; and 21 failures from 10 up-selected fish. Out of 639 data points from the test phase, there were 21 failures from 5 down-selected fish and 19 failures from 4 up-selected fish.

We used the open-access software R version 3.6.3. (R Core Team, 2020) to run all statistical analyses and generate the figures. We first tested whether selection for telencephalon size (up-selected vs down-selected lines) affected male guppies’ performance in the detour task. Then we tested whether the individual telencephalon size proportion predicted performance in a similar manner as selection lines. Furthermore, we investigated differences in the relative telencephalon size between the sampled up’ and down-selected fish. In all mixed effect models we ran here, fish identity was fitted as the random variable.

To test performance during the pre-training phase, we fitted a generalised linear mixed effect model (GLMER) with the number of time-bins until a fish reached the food reward as the response variable (using a negative binomial error distribution), selection line as categorical predictor, trial number as continuous predictor and replicate as a covariate. In addition, a similar model was fitted, but with telencephalon size proportion as a continuous predictor along with trial number.

For the test phase, we tested the time elapsed until a fish reached the food reward in a linear mixed effect model (LMER). Again, we fitted selection line as a categorical predictor, trial number as a continuous predictor, and replicate as a covariate. For testing the predictions by individual telencephalon size, a similar model was fitted with telencephalon size proportion as a continuous predictor along with trial number. The same logic applied for binomial GLMERs testing correct detours during test trials (i.e., whether fish touched the plastic cup while detouring) where the first model tested the effect of selection line, while the other model tested the effect of individual measurements of telencephalon size proportions.

Fitted models were checked for normality of residuals, homoscedasticity of the variance, and potential overdispersion or underdispersion, accordingly. For the LMER models, the variable “time” was log-transformed to meet the model’s assumptions. For every model, we also estimated the proportion of explained variance by fixed and random factors (conditional R^2^) and by fixed factors only (marginal R^2^). For *post hoc* analyses, we used functions from the Estimated Marginal Means package (`emmeans`) in R (Lenth & Lenth, 2018). This package allows *post hoc* analyses in models involving interactions between categorical factors and continuous predictors.

## Acknowledgements

We kindly thank Annika Boussard and Alberto Corral-Lopez for their help and technical advice regarding data collection.

## Ethics

This work was approved by the ethics research committee of the Stockholm Animal Research Ethical Permit Board [permit number: 17362-2019].

## Data and code

The data and code to reproduce the statistical outcomes and generate the figures are available in the repository Figshare (Data DOI: https://doi.org/10.6084/m9.figshare.12234725).

***For review purpose, the files can be accessed via a private link:*** https://figshare.com/s/a233e992b2912c4fd3ed

## Author contribution

ZT conceived and designed the study, carried out the laboratory experiment, analysed the data, and drafted the manuscript; SF conceived and designed the study and helped with data collection; MA collected the brain measurements data; NK conceived and designed the study, coordinated the study and helped draft the manuscript. ZT and NK finalised the manuscript with input from all authors. All authors gave final approval for publication and agreed to be held accountable for the work performed therein.

## References

Ashton BJ, Ridley AR, Edwards EK, Thornton A. 2018. Cognitive performance is linked to group size and affects fitness in Australian magpies. Nature 554:364–367. DOI: 10.1038/nature25503.

Barton RA, Harvey PH. 2000. Mosaic evolution of brain structure in mammals. 405:4.

Benson-Amram S, Dantzer B, Stricker G, Swanson EM, Holekamp KE. 2016. Brain size predicts problem-solving ability in mammalian carnivores. Proceedings of the National Academy of Sciences 113:2532–2537. DOI: 10.1073/pnas.l505913113.

Boogert NJ, Anderson RC, Peters S, Searcy WA, Nowicki S. 2011. Song repertoire size in male song sparrows correlates with detour reaching, but not with other cognitive measures. Animal Behaviour 81:1209–1216. DOI: 10.1016/j.anbehav.2011.03.004.

Boogert NJ, Madden JR, Morand-Ferron J, Thornton A. 2018. Measuring and understanding individual differences in cognition. Philosophical Transactions of the Royal Society B: Biological Sciences 373:20170280. DOI: 10.1098/rstb.2017.0280.

Brandão, ML, Fernandes AMT de A, Gonçalves-de-Freitas E. 2019. Male and female cichlid fish show cognitive inhibitory control ability. Scientific Reports 9:15795. DOI: 10.1038/s41598-019-52384-2.

Briscoe SD, Ragsdale CW. 2019. Evolution of the Chordate Telencephalon. Current Biology 29:R647–R662. DOI: 10.1016/j.cub.2019.05.026.

Buechel SD, Boussard A, Kotrschal A, van der Bijl W, Kolm N. 2018. Brain size affects performance in a reversal-learning test. Proceedings of the Royal Society B: Biological Sciences 285:20172031. DOI: 10.1098/rspb.2017.2031.

Costa SS, Andrade R, Carneiro LA, Gonçalves EJ, Kotrschal K, Oliveira RF. 2011. Sex differences in the dorsolateral telencephalon correlate with home range size in blenniid fish. Brain, Behavior and Evolution 77:55–64. DOI: 10.1159/000323668.

Deaner RO, Isler K, Burkart J, van Schaik C. 2007. Overall brain size, and not encephalization quotient, best predicts cognitive ability across non-human primates. Brain, Behavior and Evolution 70:115–124. DOI: 10.1159/000102973.

DeCasien AR, Higham JP. 2019. Primate mosaic brain evolution reflects selection on sensory and cognitive specialization. Nature Ecology & Evolution 3:1483–1493. DOI: 10.1038/s41559-019-0969-0.

Ebbesson LOE, Braithwaite VA. 2012. Environmental effects on fish neural plasticity and cognition. Journal of Fish Biology 81:2151–2174. DOI: 10.1111/j.1095-8649.2012.03486.x.

Fagnani J, Barrera G, Carballo F, Bentosela M. 2016. Is previous experience important for inhibitory control? A comparison between shelter and pet dogs in A-not-B and cylinder tasks. Animal Cognition 19:1165–1172. DOI: 10.1007/s10071-016-1024-z.

Finlay BL, Darlington RB, Nicastro N. 2001. Developmental structure in brain evolution. Behavioral and Brain Sciences 24. DOI: 10.1017/S0140525X01423958.

Fong S, Rogell B, Amcoff M, Kotrschal A, Bijl W van der, Buechel SD, Kolm N. 2021. Rapid mosaic brain evolution under artificial selection for relative telencephalon size in the guppy (Poecilia reticulata). Preprint, bioRxiv:2021.03.31.437806. DOI: 10.1101/2021.03.31.437806.

Gonzalez-Voyer A, Kolm N. 2010. Sex, Ecology and the Brain: Evolutionary Correlates of Brain Structure Volumes in Tanganyikan Cichlids. PLOS ONE 5:e14355. DOI: 10.1371/journal.pone.0014355.

Gonzalez-Voyer A, Winberg S, Kolm N. 2009. Brain structure evolution in a basal vertebrate clade: evidence from phylogenetic comparative analysis of cichlid fishes. BMC Evolutionary Biology 9:238. DOI: 10.1186/1471-2148-9-238.

Jerison H. 1973. Evolution of the Brain and Intelligence.New York: Academic Press New York: Academic Press.

Juszczak GR, Miller M. 2016. Detour behavior of mice trained with transparent, semitransparent and opaque barriers. PloS one 11:e0162018.

Kabadayi C, Bobrowicz K, Osvath M. 2018. The detour paradigm in animal cognition. Animal Cognition 21:21–35. DOI: 10.1007/s10071-017-1152-0.

Kolm N, Gonzalez-Voyer A, Brelin D, Winberg S. 2009. Evidence for small scale variation in the vertebrate brain: mating strategy and sex affect brain size and structure in wild brown trout (Salmo trutta). Journal of Evolutionary Biology 22:2524–2531. DOI: 10.1111/j.l420-9101.2009.01875.x.

Kotrschal A, Rogell B, Bundsen A, Svensson B, Zajitschek S, Brännström I, Immler S, Maklakov AA, Kolm N. 2013. Artificial Selection on Relative Brain Size in the Guppy Reveals Costs and Benefits of Evolving a Larger Brain. Current Biology 23:168–171. DOI: 10.1016/j.cub.2012.11.058.

Lenth R, Lenth MR. 2018. Package ‘lsmeans.’ The American Statistician 34:216–221.

López JC, Bingman VP, Rodríguez F, Gómez Y, Salas C. 2000. Dissociation of place and cue learning by telencephalic ablation in goldfish. Behavioral Neuroscience 114:687–699.

Lucon-Xiccato T, Gatto E, Bisazza A. 2017. Fish perform like mammals and birds in inhibitory motor control tasks. Scientific Reports 7:13144. DOI: 10.1038/s41598-017-13447-4.

MacLean EL, Hare B, Nunn CL, Addessi E, Amici F, Anderson RC, Aureli F, Baker JM, Bania AE, Barnard AM, Boogert NJ, Brannon EM, Bray EE, Bray J, Brent UN, Burkart JM, Call J, Cantlon JF, Cheke LG, Clayton NS, Delgado MM, DiVincenti LJ, Fujita K, Herrmann E, Hiramatsu C, Jacobs LF, Jordan KE, Laude JR, Leimgruber KL, Messer EJE, Moura AC de A, Ostojić L, Picard A, Platt ML, Plotnik JM, Range F, Reader SM, Reddy RB, Sandel AA, Santos LR, Schumann K, Seed AM, Sewall KB, Shaw RC, Slocombe KE, Su Y, Takimoto A, Tan J, Tao R, Schaik CP van, Virányi Z, Visalberghi E, Wade JC, Watanabe A, Widness J, Young JK, Zentall TR, Zhao Y. 2014. The evolution of self-control. Proceedings of the National Academy of Sciences 111:E2140–E2148. DOI:10.1073/pnas.1323533 111.

Miyake A, Friedman NP, Emerson MJ, Witzki AH, Howerter A, Wager TD. 2000. The unity and diversity of executive functions and their contributions to complex “frontal lobe” tasks: A latent variable analysis. Cognitive psychology 41:49–100.

O’Connell, Hofmann HA. 2011. The Vertebrate mesolimbic reward system and social behavior network: A comparative synthesis. The Journal of Comparative Neurology 519:3599–3639. DOI: 10.1002/cne.22735.

Pollen AA, Dobberfuhl AP, Scace J, Igulu MM, Renn SCP, Shumway CA, Hofmann HA. 2007. Environmental Complexity and Social Organization Sculpt the Brain in Lake Tanganyikan Cichlid Fish. Brain, Behavior and Evolution 70:21–39. DOI: 10.1159/000101067.

Portavella M, Vargas JP, Torres B, Salas C. 2002. The effects of telencephalic pallial lesions on spatial, temporal, and emotional learning in goldfish. Brain Research Bulletin 57:397–399. DOI: 10.1016/S0361-9230(01)00699-2.

R Core Team. 2020. A language and environment for statistical computing. Vienna, Austria: R Foundation for Statistical Computing.

Reader SM, Hager Y, Laland KN. 2011. The evolution of primate general and cultural intelligence. Philosophical Transactions of the Royal Society B: Biological Sciences 366:1017–1027. DOI: 10.1098/rstb.2010.0342.

Salas C, Broglio C, Rodríguez F, López JC, Portavella M, Torres B. 1996. Telencephalic ablation in goldfish impairs performance in a ‘spatial constancy’ problem but not in a cued one. Behavioural Brain Research 79:193–200. DOI: 10.1016/0166-4328(96)00014-9.

Santos LR, Ericson BN, Hauser MD. 1999. Constraints on problem solving and inhibition: Object retrieval in cotton-top tamarins (Saguinus oedipus oedipus). Journal of Comparative Psychology 113:186.

van Schaik CP, Isler K, Burkart JM. 2012. Explaining brain size variation: from social to cultural brain. Trends in Cognitive Sciences 16:277–284. DOI: 10.1016/j.tics.2012.04.004.

Shettleworth SJ. 2010. Cognition, Evolution, and Behavior. Oxford University Press.

Shultz S, Dunbar RIM. 2010. Species differences in executive function correlate with hippocampus volume and neocortex ratio across nonhuman primates. Journal of Comparative Psychology 124:252–260. DOI: 10.1037/a0018894.

Striedter GF. 2005. Principles of brain evolution. USA: Sinauer Associates.

Striedter GF, Northcutt RG. 2019. Brains Through Time: A Natural History of Vertebrates. Oxford University Press.

Taborsky B, Oliveira RF. 2012. Social competence: an evolutionary approach. Trends in Ecology & Evolution 27:679–688. DOI: 10.1016/j.tree.2012.09.003.

Thornton A, Lukas D. 2012. Individual variation in cognitive performance: developmental and evolutionary perspectives. Philosophical Transactions of the Royal Society B: Biological Sciences 367:2773–2783. DOI: 10.1098/rstb.2012.0214.

Timmermans S, Lefebvre L, Boire D, Basu P. 2000. Relative Size of the Hyperstriatum ventrale Is the Best Predictor of Feeding Innovation Rate in Birds. Brain, Behavior and Evolution 56:196–203. DOI: 10.1159/000047204.

Triki Z, Emery Y, Teles MC, Oliveira RF, Bshary R. 2020. Brain morphology predicts social intelligence in wild cleaner fish. Nature Communications 11:6423. DOI: 10.1038/s41467-020-20130-2.

Triki Z, Levorato E, McNeely W, Marshall J, Bshary R. 2019a. Population densities predict forebrain size variation in the cleaner fish *Labroides dimidiatus*. Proceedings of the Royal Society B: Biological Sciences 286:20192108. DOI: 10.1098/rspb.2019.2108.

Triki Z, Wismer S, Rey O, Ann Binning S, Levorato E, Bshary R. 2019b. Biological market effects predict cleaner fish strategic sophistication. Behavioral Ecology 30:1548–1557. DOI: 10.1093/beheco/arzlll.

Wallis JD, Dias R, Robbins TW, Roberts AC. 2001. Dissociable contributions of the orbitofrontal and lateral prefrontal cortex of the marmoset to performance on a detour reaching task. European Journal of Neuroscience 13:1797–1808. DOI: 10.1046/j.0953-816x.2001.01546.x.

White GE, Brown C. 2015. Microhabitat Use Affects Brain Size and Structure in Intertidal Gobies. Brain, Behavior and Evolution 85:107–116. DOI: 10.1159/000380875.

